# Detection of hypermucoviscous *Klebsiella pneumoniae* sequence type 86 capsular type K2 in South America as an unexpected cause of a fatal outbreak in captivity marmosets

**DOI:** 10.1101/2020.02.02.930685

**Authors:** Juliana M. Guerra, Natália. C.C. de A. Fernandes, Alessandra Loureiro Morales dos Santos, Joana de Souza Pereira Barrel, Bruno Simões Sergio Petri, Liliane Milanelo, Monique Ribeiro Tiba-Casas, Alcina Maria Liserre, Cláudia Regina Gonçalves, Cláudio Tavares Sacchi, José Luiz Catão-Dias, Carlos Henrique Camargo

## Abstract

After the sudden death of eleven captives marmosets in a rehabilitation center of wildlife in São Paulo, Brazil, histological and microbiological study was conducted. Liver, spleen, intestine, central nervous system, lung, thymus, stomach, testicle tissues were analyzed by light microscopy and microbial cultures were conducted. Environmental cultures were also performed. Prophylactic antimicrobial therapy, restricted access to marmosets’ cages with dedicated staff, and additional sanitization of animals’ fruits were implemented. Histological findings were compatible with hyperacute septicemia, and microbiological cultures and molecular tests identified the etiologic agent as hypermucoviscous sequence type 86 capsular type K2 *K. pneumoniae* for the first time in South America. Implementation of prompt containment measures led to successful control of this outbreak. Detection of a hypervirulent and zoonotic pathogen, such as hypermucoviscous *K. pneumoniae* ST86 K2, in an unexpected and human interface reservoir underscores its potential threat in public health settings.

## 1. INTRODUCTION

*Klebsiella pneumoniae* is a Gram-negative, encapsulated, non-motile, rod-shaped and opportunistic bacteria present as a normal part of the nasopharynx and gastrointestinal tract microbiome of humans and animals (1–3). Since the mid-1980s and 1990s, a new hypervirulent (hypermucoviscous) variant of *K. pneumoniae* (hvKp), initially described in Southeastern Asia (4–6), has arisen as an important emerging pathogen, affecting young and healthy individuals from Europe, North, Central and South America, Australia, Africa and the Middle East (7–13).

This variant presents clinical signs and bacterial genotypic and phenotypic features that allow the differentiation from opportunistic *K. pneumoniae*. hvKp has been frequently associated with serious clinical complications, including multisystemic pyogenic abscess in liver, lungs and brain, meningitis, endophthalmitis, osteomyelitis, septicemia and pulmonary embolism (12–15). Also, hvKp strains usually have a distinguish hypermucoviscosity phenotype when grown on agar plates, based on a positive string test. The development of prominent polysaccharide capsules, associated with capsular serotypes K1 or K2, have been reported as the major virulence determinants for human hvKp in liver abscesses (16–19), since its seems to protect the bacteria from phagocytosis and preventing destruction by bactericidal serum factors. In addition, a number of putative virulence genes, such as the gene *rmpA*, a regulator of the mucoid phenotype gene, located on a plasmid, mucoviscosity-associated gene *magA*, which encodes a structural outer membrane protein of the K1 serotype, and aerobactin genes have been described (17,20,21).

*K. pneumoniae* strains have also been associated with a variety of diseases in animals, especially in New and Old World primates (22–24). Sudden death or variable clinical signs, including anorexia, prostration, fever, cough, dyspnea, mucopurulent discharge, meningitis, pneumonia, peritonitis, and sepsis are strongly associated with sporadic infections of *Klebsiella pneumoniae* in experimental colonies of common marmosets (24,25). A mixed infection of *Klebsiella* species and *Bordetella bronchiseptica* have also been reported to cause purulent otitis media in this same species, with frequent extension to the brain, leading to abscessing encephalitis and meningitis (26). Multiple antibiotic-resistant *Klebsiella* strains have been isolated from meningitis in *Macaca mulatta* (rhesus macaques) (22). Richard (1989) described a subcutaneous abscess due to *K. pneumoniae* K5 in a breed of squirrel monkey (*Saimiri* spp.) at the Pasteur Institute of Cayenne, French Guyana, and a fatal infection due to *K. pneumoniae* K2 in a colony of lemurs, at Mulhouse zoological garden (East of France) (27). Positive *K. pneumoniae* bacterial culture related to necrotic and hemorrhagic enteritis and typhlocolitis were observed in an experimental colony of owl monkeys (*Aotus nancymai*) (2). More recently, hvKp has been identified in *Cercopithecus aethiops* (Africa green monkeys) causing liver abscesses with bacteremia, meningitis, endophthalmitis (28–30) and suppurative in a captive gold‐handed tamarin (*Saguinus midas midas*) (31). Bilateral mucopurulent nasal discharge, fever, anemia, and swelling of the inguinal lymph nodes was described in a *Allouatta Clamitans* due to hvKp ST23 serotype K1 (32). However, some species, as cynomolgus and rhesus macaques can act as carriers and maintain subclinical infection with *K. pneumoniae* (30).

Despite the well-recognized zoonotic importance of hvKp and the public health risk of emerging multidrug-resistant strains (33–35), there is a lack of information regarding the complete genotyping and phenotyping characterization of the etiologic agent, which is essential to establish adequate diagnosis and treatment of this pathogen in captive and wild non-human primates. Thus, the aim of this study was to report an epizootic of common marmosets in a rehabilitation center of wildlife in Brazil and characterize the serotype, sequence typing, virulence properties and resistance profile of *K. pneumoniae* strains.

## 2. MATERIALS AND METHODS

### 2.1 Study population

All the animals described in this study were maintained in Parque Ecológico do Tietê (23°29’46“S, 46°31’10“O), a center for reception, rehabilitation and referring of wildlife animals, located at São Paulo municipality, São Paulo, Brazil. All callitrichids (*Callithrix jacchus, C. penicillata* and hybrids) were housed in a shed of five stainless steel 2 m³ cages and 15 stainless steel 0.4 m³ cages, dedicated only to this species, with a maximum number of eight animals per cage with anti-mosquito screen, feeding platforms, logs and ropes. This center comprises an average daily census of approximately 70 wild-caught callitrichids, that were brought to the facility due to hunting or traffic. These animals have been maintained in captivity for an average period of 140 days for rehabilitation. The cages were cleaned daily by an exclusive employee, except on weekends, when the same worker also cleans the reptilian cages. Standard husbandry procedures included two-times daily feeding program with a commercial food (Megazoo, Minas Gerais, Brazil), diet supplements consisting of a variety of vegetables, fruits, sporadically, *Tenebrio* spp. and crickets, and water *ad libitum*. Routine quarantine procedures consist of physical examination and vermifugation. The animals were also periodically monitored by a veterinarian.

### 2.2 Animals and sampling

On 10th and 11 February 2019, eleven captive marmosets (eight *C. penicillata*, two C. *jacchus* and one hybrid), without previous clinical signs, suddenly died. They lived in the same shed and into five different cages with others 72 animals and none of them presented any symptoms. All animals were in captivity from 123 to 399 days. No new animals had been introduced into the cages in the last 25 days.

The necropsy was performed on the same day and macroscopic examination showed no major findings. Tissues samples were preserved in phosphate-buffer formalin 10% and in refrigeration for 12 hours. Since 1999, according to the Non-Human Primates Epizootic Events Surveillance System of Brazilian Ministry of Health (36), which was incorporated into the Yellow Fever Surveillance System. All epizootic event in this population should be investigated in reference laboratories. As so, these samples were sent to Adolfo Lutz Institute for analysis. This study was approved by the Ethics Committee for the Use of Animals (CEUA) of Adolfo Lutz Institute, Brazil (protocol n°11/2016), SISBIO registration no. 50551 for the manipulation of wildlife material and SISGEN registration no. A1A2A72 and A7EB4B6.

### 2.3 Histological and histochemical examination

For light microscopy small pieces of liver, spleen, intestine, central nervous system, lung, thymus, stomach, and testicle tissue were fixed in neutral 10% phosphate-buffered saline formaldehyde. Fixed tissue specimens were dehydrated by graded ethanol treatment and were routinely embedded in paraffin wax for light microscopy. Embedded sections (3 μm) were stained with haematoxylin and eosin (H&E). Also, additional slides were submitted to histological gram staining. The presence of lesions, intensity, distribution and presence of inflammatory infiltrate were analyzed.

### 2.4 Isolation and characterization of *Klebsiella* spp

Liver, brain and pooled tissues were aseptically cultured on sheep blood agar (Oxoid, Basingstoke, UK) and MacConkey agar (Oxoid, Basingstoke, UK) and incubated overnight at 37°C. The plates were examined the day after inoculation for the presence of any growth. Pink-yellow, mucoid, lactose-positive colonies were cultured, and identified as *Klebsiella pneumoniae* Complex based on motility, lysine decarboxylation, citrate utilization and indole non-production. The string test was used to check the hypermucoviscosity of *K. pneumoniae* colonies. The result was defined as positive when an inoculation loop was able to generate a viscous string of 5 mm in length from a colony on a blood agar plate after overnight incubation (8) while a negative sting test was assigned when the string was less than 5 mm in length or no string was present.

### 2.5 Environmental testing

In order to identify the source of *K. pneumoniae* strains, environmental samples of water from the lake near to the primate cages and drag swabs from their cages were screened for the presence of *K. pneumoniae.* Water samples (100mL) were filtered with 0.22 μm sterile membranes (Jet Biofil, China) and placed onto the surface of a MacConkey Agar plate. Typical colonies (mucoid, lactose-fermenting) were identified by biochemical tests (see above). Drag swabs were streaked onto MacConkey Agar plates and typical *K. pneumoniae* colonies were subjected to the same tests as employed for the water isolates.

### 2.6 Antimicrobial susceptibility testing

A representative isolate (P04) was subjected to antimicrobial susceptibility testing by using the broth microdilution methodology with Sensititre (Thermo, Waltham, USA) plate, according to the manufacturer’s instructions. Minimal inhibitory concentration values were categorized as susceptible, intermediate or resistance following Clinical and Laboratory Standards Institute (CLSI, Wayne, USA) M100-S29 breakpoints (available at http://em100.edaptivedocs.net/dashboard.aspx).

### 2.7 Pulsed-field gel electrophoresis (PFGE)

All isolates identified as *K. pneumoniae* were submitted to DNA macro-restriction by using 30U of XbaI enzyme followed by Pulsed-field gel electrophoresis in a CHEF-III apparatus, according to the PulseNet International guidelines (https://www.cdc.gov/pulsenet/pathogens/pfge.html). The Universal Size Standard Strain H9812 (*Salmonella* Braenderup) (37) was used as reference in all gels. Gel images were analyzed in the BioNumerics v.7.6.2 (Applied Maths, Sint-Martens-Latem, Belgium) with the generation of dendrogram based on unweighted pair group method with arithmetic mean (UPGMA) methods and similarities calculated by Dice coefficient with tolerance and optimization both set at 1.5%.

### 2.8 DNA extraction and whole genome sequencing

The representative strain P04 was selected based on the PFGE results, and subjected to whole genome sequencing. High quality whole genome was extracted from a pure culture of 16h growth in Luria-Bertani broth (Oxoid, Basingstoke, UK) by using the Wizard Genomic DNA Purification Kit (Promega, Madison, USA) according to the instructions of the manufacturer. DNA was tagmented by using the Ion Xpress™ Plus Fragment Library Kit and the libraries constructed were sequenced in an Ion Torrent S5 platform. Reads were de novo assembled with the Spades algorithm (v.5.12.0.0) and submitted to online platforms. Definitive species identification was carried out by Ribosomal Multilocus Sequence Typing (rMLST) (https://pubmlst.org/rmlst/). Antimicrobial resistance genes were sought at the Resfinder tool of the Center for Genomic Epidemiology website (http://www.genomicepidemiology.org/). Sequence Type, virulence genes and capsular type data were extracted from the Institute Pasteur MLST and whole genome MLST databases (http://bigsdb.pasteur.fr/). Based on the reference genome CG43 (isolate 1530 on Institute Pasteur database available at https://bigsdb.pasteur.fr/cgi-bin/bigsdb/bigsdb.pl?db=pubmlst_klebsiella_isolates&page=plugin&name=Contigs&format=text&isolate_id=1530&match=1&pc_untagged=0&min_length=&header=1), high quality single-nucleotide polymorphisms (SNPs) were extracted from the P04 genome and from the available ST86 *K. pneumoniae* genomes from the Institute Pasteur MLST and whole genome MLST databases. A phylogenetic tree based on the concatenated alignments was built on the CSI Phylogeny 1.4 (https://cge.cbs.dtu.dk/services/CSIPhylogeny/). The genome of the hv ST374 *K. pneumoniae* was included as outgroup. The Newick generated file was uploaded on the Microreact platform along with metadata of such isolates (project V8uV0aPEU) for the generation of a tree with the SNPs and metadata. The whole-genome shotgun project reported here has been deposited at DDBJ/EMBL/GenBank under the accession number SPSP00000000.

## RESULTS

### 2.1 Histological and histochemical examination

Multiple sections of different tissues were microscopically examined and similarly affected. The major microscopic finding consistent to sepsis it was intravascular presence of bacilli in liver (10/10), cerebrum (8/10), lung (3/10), heart (1/7), intestine (1/7), thymus (1/2), and skeletal muscle (1/1). Others relevant microscopic findings of different tissues are summarized on the Table 1.

Furthermore, intestine it was the unique organ that it was observed bacteria into the lumen of lymphatic vessels (1/7). Other nonspecific findings consists to tubular proteinosis (4/8) and tubular acute necrosis (1/8) in kidney, hepatic steatosis (1/10), lymph histiocytic admixed or not with neutrophils enteritis (2/7), Peyer plaques hyperplasia (1/7), cardiac congestion (1/7), focus of edema of the cardiomyocytes and necrotizing and lymphohistiocytic myocarditis (1/7). Stomach, tongue, testis, thymus, skin, uterus, lymph-node didn’t show microscopic alterations on all the fragments analyzed. Some tissues couldn’t be analyzed due to the several autolysis.

### 2.2 Isolation and characterization of *Klebsiella* spp

Microbial cultures performed from brain and liver from eight different animals recovered *K. pneumoniae* in all of them, as pure growth, and with hypermucoviscous phenotype, i.e., positive string test. The isolate P04 presented susceptibility to all the antimicrobial agents evaluated (Table 2).

### 2.3 Environmental testing

*K. pneumoniae* isolates were recovered from both water and drag swab samples.

### 2.4 Pulsed-field gel electrophoresis (PFGE)

Molecular typing of *K. pneumoniae* revealed the same XbaI-PFGE profile among the isolates recovered from the dead animals (Figure 2, cluster A, pulsotypes A1 and A2), however, environmental isolates clustered apart from the invasive isolates (Figure 2, clusters B and C).

**Figure 1.**
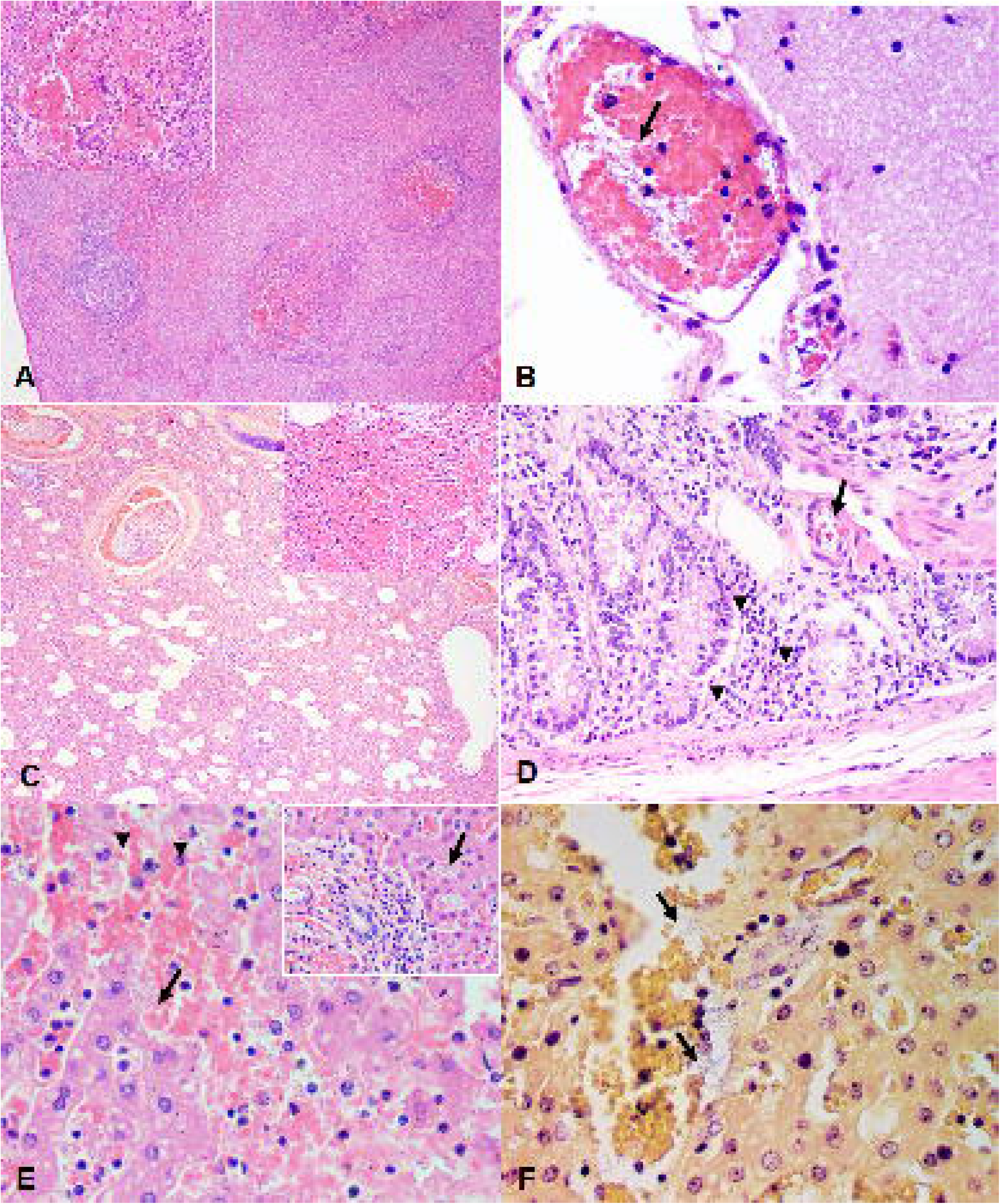
Microscopic findings of histological and histochemical examination prepared for light microscopy. A – Spleen. Necro hemorrhagic splenitis in multiple germ centers (H&E 4x). B – Brain (Meningis). Bacterial rods inside vascular lumen (arrows) (H&E 40x). C – Lung. Several interstitial pneumonia (H&E 4x) and alveolar hemorrhage (inset – H&E10x). D – Intestine. Lymphoplasmacytic enteritis (arrowheads) and intravascular bacterias (arrow) (H&E 20x and 40x). E – Liver. Necrosis (arrowheads) associated with the presence of numerous bacterial rods (arrow). Hepatocytes also show steatosis and the sinusoids are filled by neutrophils (H&E 40x). The inset image shows discrete mononuclear portal hepatitis and presence of numerous bacterial rods (H&E 20x). F – Liver. The sinusoids are filled by gram-negative bacterial structures (heads) and neutrophils (Gram 40x).

**Figure 2.**
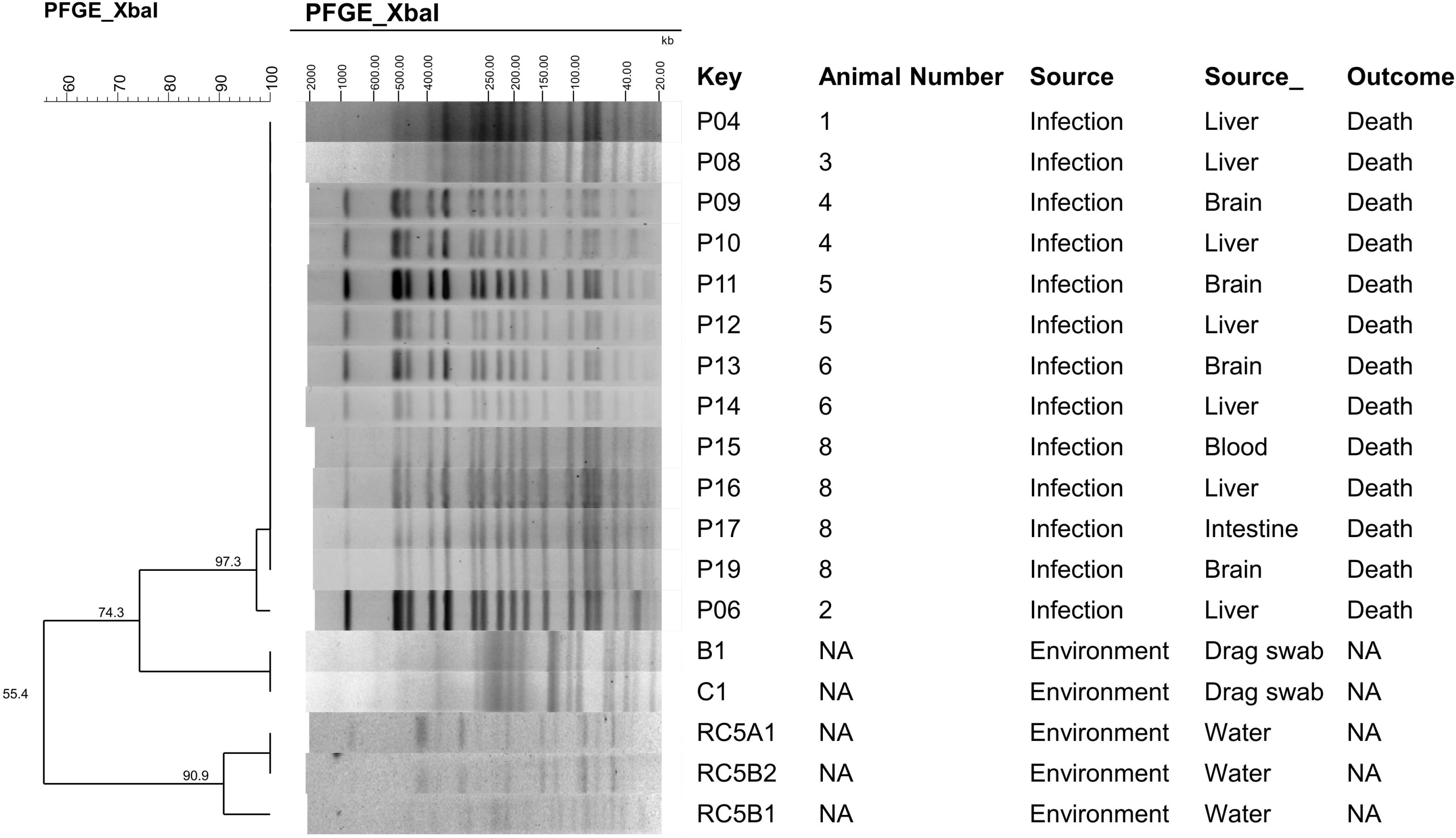
Dendrogram and PFGE typing of XbaI restricted *K. pneumoniae* strains isolated from infection (blue, cluster A), drag swab (green, cluster B) and water (red, cluster C).PFGE profiles (represented by capital letters) were defined based on 100% Dice similarity cutoff value of the UPGMA clustering method (1.5% optimization; 1.5% tolerance). NA: not applicable.

### 2.5 Whole genome sequencing

Whole genome sequencing resulted in 1,049,163 reads and the *de novo* assembled genome comprised 5,358,608 bps grouped in 62 contigs, with an average coverage depth of 53. rMLST and kmer tools confirmed the presence of *K. pneumoniae* as the species of isolate P04. Acquired antimicrobial resistant genes were not detected, but the constitutive *bla*SHV-1, *oqx*AB and *fos*A were. *In silico* multilocus sequence typing analysis identified the isolate as the sequence type ST86, with the following combination alleles: *gapA*, 9; *infB*, 4; *mdh*, 2; *pgi*, 1; *phoE*, 1; *rpoB*, 1; *tonB*, 27. The capsular type K2 was identified. According to the virulence database of BIGSdb *K. pneumoniae*, P04 presented the *mrkABCDFHIJ* cluster (mannose-resistant *Klebsiella*-like type III fimbriae cluster, associated with adhesiveness and fimbrial filament formation to adherence to eukaryotic cells) (38) in the same contig of the *kvgAS* genes. The *iroBCD* genes (salmochelin) were also detected, in the same contig that the *rmpA* gene (regulator of mucoid phenotype A). The *rmpA2* gene was detected within the contig along the *iucABD* with the *iutA* genes (aerobactin).

Phylogenetic analysis of high quality SNPs showed that ST86 isolates were closely related, and the P04 strain clustered close to IPEUC340, an isolate recovered in 1975 in France (Figure 3.A). The isolate RJF293, ST374, clustered apart from the ST86 isolates (Figure 3.B).

**Figure 3.**
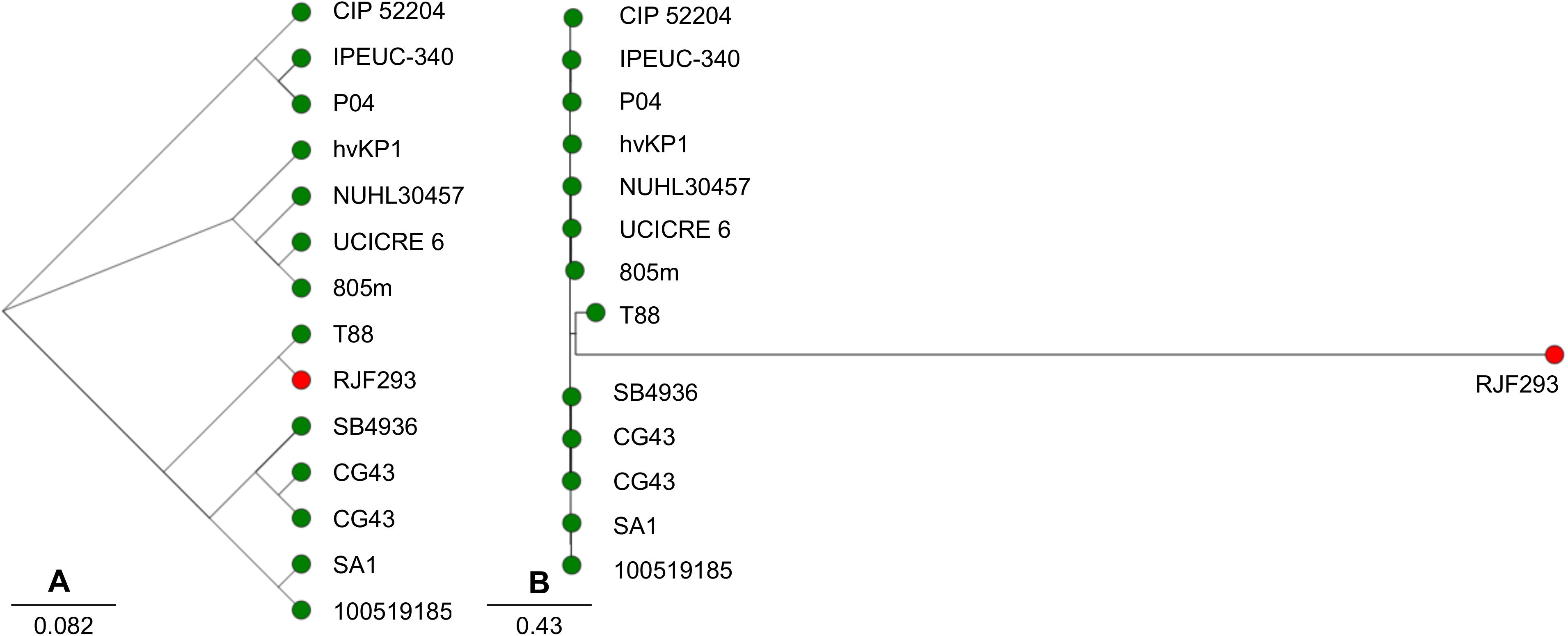
Schematic representations of high quality SNPs trees built from publicly available ST86 *K. pneumoniae* genomes. In (A) it is possible to note that the P04 isolate described in this study clustered close to the IPEUC-340 isolate, recovered in 1975 in France. In (B) we observe the high similarity of ST86 *K. pneumoniae* genomes in comparison with the outgroup strain RJF293, an ST374 hypermucoviscous *K. pneumoniae* recovered from invasive infection. Green circles represent the ST86 isolates, and the red one represents the ST374.

## DISCUSSION

In this study, eleven marmosets suddenly died after habitual management of the captives in one of the largest centers for reception, rehabilitation and referring of wildlife animals in the municipality of São Paulo, the largest city in South America. Histological findings were compatible with a hyperacute septicemia, microbiological cultures and molecular tests identified the etiologic agent as hypermucoviscous, ST86, K2 *K. pneumoniae.* This bacteria represents one of the priority organisms to be controlled according to the World Health Organization, particularly the carbapenem-and polymyxin-resistant strains (39). The emergence of these strains is a concern in human and veterinary medicine due to the potential of these strains to acquire multidrug resistance genes, their capacity to persist in the environment and to infect a wide-range of hosts and the unavailability of vaccines against these strains for humans and animals (40).

The role of *K. pneumoniae* in wildlife has been revisited, especially in callitrichidae species, which are highly susceptible to fatal infections caused by this bacteria in captivity (24,40). The bacteria likely spreads by close contact and aerosol between animals and through fomites between cages (41). Frequently described in association with immunosuppressive conditions in New world monkeys, *Klebsiella* is considered an opportunist agent, leading to pneumonia, peritonitis, cystitis, meningitis, and septicemia (2). The propensity of common marmosets to bacterial infection could be related to the low variability in the MHC class II loci that may compromise anti-bacterial immune response and also explain the high frequency of septicemia and endotoxemia after deep wounds and fighters in this genus (41,42).

We identified that the strains recovered from marmosets that suddenly died were clonally related by PFGE analysis. Since PFGE is known by its high discriminatory power, we can suppose that a common source was present. In order to investigate the source of such strain, environmental samples (water and drag swabs) were cultured, and despite the growth of *K. pneumoniae*, molecular typing revealed that environmental strains were not related to the clinical isolates. Despite our efforts, we were not able to detect the source of the hypermucoviscous *K. pneumoniae* in this outbreak during our investigation. Other studies also faced this same limitation (2) and suggested that dietary supplements could be the most likely source responsible for the introduction of *K. pneumoniae* in the colonies. Vegetables and fruits, especially bananas and their peels, seems to carry a widely diverse Enterobacteriales microbial community (23). Even we have not found the source of the virulent *K. pneumoniae* isolate causing this outbreak, drag swabs of the living animals were negative for such strain, indicating that the remaining population of marmosets was not carriers of this hvKp isolate. In fact, no other animals died after this event. We can admit that the end of this outbreak can be attributed to implementation of effective measures to contain the dissemination of the virulent *K. pneumoniae* strain. The use of Bactrim as prophylaxis during the first 5 days after the identification of deaths may be one of the successful measures implemented, since the isolate was susceptible to this drug. In fact, the isolate did not present any resistance marker to the evaluated antimicrobial agents (Table 2). In addition, accession to the cages was restricted to a dedicated employee, who wore dedicated clothes during the permanence in the marmosets’ shed. Additional sanitization step of fruits with 2% sodium hypochlorite for 15 minutes and dedicated space and staff for preparation of marmosets’ meals were also implemented after the epizootic event.

Whole genome sequencing of one representative isolate identified several virulence genes that can be associated with its hypermucoviscous phenotype. Among the several virulence genes detected, the ones associated with invasiveness and with capsule production could contribute to the evasiveness of immune system resulting in the severe and fulminant clinical conditions observed. Indeed, four out of the five suggested virulence genes recognized to be associated with the hv phenotype were detected in the P04 genome, namely *rmpA, rmpA2* (regulators of mucoid phenotype A), *iuc* (aerobactin), and *iroB* (salmochelin) (43).

The hypermucoviscous *K. pneumoniae* P04 strain was identified as sequence type ST86 and capsular typing K2. Previous reports also identified this hypervirulent clone with a wild-type susceptibility profile causing severe infections in both hospital (44) and community-acquired infections in humans (45), highlighting the role of hypermucoviscous *K. pneumoniae* as etiological of human infections. Intriguingly, infections due hvKp ST86 K2 are apparently more severe than the infections due to other hvKp clones (46). As of October 2019, 26 isolates belonging to ST86 were available in the PasteurMLST database (https://bigsdb.pasteur.fr/). All of them isolated from humans, and predominantly from Europe (12/26) and Asia (10/26) (10,47–49). Only two isolates were previously identified in America, one recovered from the blood of a patient with liver abscess in Buffalo, USA, in 2013 (50) and another one recovered from the cerebrospinal fluid of a patient with meningitis in Pointe-a-Pitre, Guadeloupe, in 2014 (51). Therefore, to the best of our knowledge, this is the first description of hvKp ST86 K2 *K. pneumoniae* from South America.

Our phylogenetic analysis demonstrated that ST86 genomes are very closely related, but the origin of the P04 isolate could not be certainly inferred, since it clustered with the RFJ293 isolate, recovered in 1975 from France. The expansion of the emerging pathogen hypermucoviscous *K. pneumoniae* ST86 K2 among different reservoirs should be carefully surveilled, since relates of hypervirulent and multi-drug resistant strains are raising (52).

In summary, we report the detection of hypermucoviscous *K. pneumoniae* ST86 K2 for the first time in South America, responsible for a sudden and fatal outbreak in captive marmosets. Implementation of prompt containment measures led to successful control of this outbreak. The burden of hypermucoviscous *K. pneumoniae* ST86 K2 in unexpected reservoirs, including the ones in human interface, deserves further investigation.

## ACKNOWLEDGEMENTS

We thank the team of curators of the Institut Pasteur MLST and whole genome MLST databases for curating the data and making them publicly available at http://bigsdb.pasteur.fr/ and all contributors of Center of Pathology for routine sample processing.

## CONFLICT OF INTEREST

The authors declare that they have no conflict of interest.

## Tables headings

Table 1. Histologic findings described about the fragments submitted for microscopic evaluation.

Table 2. Antimicrobial susceptibility profile determined by Sensititre (Thermo Scientific) of the epizootic strain hypermucoviscous *K. pneumoniae* ST86 strain P04.

## REFERENCES

1. Davis GS, Price LB. Recent Research Examining Links Among Klebsiella pneumoniae from Food, Food Animals, and Human Extraintestinal Infections. Curr Environ Heal Reports. 2016 Jun;3(2):128–35.

2. Gozalo AS, Elkins WR, Lambert LE, Stock F, Thomas ML, Woodward RA. Genetic diversity of Klebsiella pneumoniae isolates during an outbreak in a non-human primate research colony. J Med Primatol. 2016 Dec;45(6):312–7.

3. Simmons J GS. Bacterial and mycotic diseases of nonhuman primates. In: Bennett BT, Abee CR HR, editor. Nonhuman Primates in Biomedical Research. San Diego: Academic Press; 2012. p. 105–72.

4. Liu YC, Cheng DL, Lin CL. Klebsiella pneumoniae liver abscess associated with septic endophthalmitis. Arch Intern Med. 1986 Oct;146(10):1913–6.

5. cheng D-L. Septic Metastatic Lesions of Pyogenic Liver Abscess. Arch Intern Med. 1991 Aug;151(8):1557.

6. Wang J, Liu Y, Lee SS, Yen M, Wang, Yao‐Shen Chen J, Wann S, et al. Primary Liver Abscess Due to Klebsiella pneumoniae in Taiwan. Clin Infect Dis. 1998 Jun;26(6):1434–8.

7. Fang FC, Sandler N, Libby SJ. Liver Abscess Caused by magA+ Klebsiella pneumoniae in North America. J Clin Microbiol. 2005 Feb;43(2):991–2.

8. Siu LK, Yeh K-M, Lin J-C, Fung C-P, Chang F-Y. Klebsiella pneumoniae liver abscess: a new invasive syndrome. Lancet Infect Dis. 2012 Nov;12(11):881–7.

9. Shon AS, Bajwa RPS, Russo TA. Hypervirulent (hypermucoviscous) Klebsiella pneumoniae. Virulence. 2013 Feb;4(2):107–18.

10. Bialek-Davenet S, Criscuolo A, Ailloud F, Passet V, Jones L, Delannoy-Vieillard A-S, et al. Genomic Definition of Hypervirulent and Multidrug-Resistant Klebsiella pneumoniae Clonal Groups. Emerg Infect Dis. 2014 Nov;20(11):1812–20.

11. Yang Z, Liu W, Cui Q, Niu W, Li H, Zhao X, et al. Prevalence and detection of Stenotrophomonas maltophilia carrying metallo-Î^2^-lactamase blaL1 in Beijing, China. Front Microbiol. 2014 Dec;5.

12. Struve C, Roe CC, Stegger M, Stahlhut SG, Hansen DS, Engelthaler DM, et al. Mapping the Evolution of Hypervirulent Klebsiella pneumoniae. Segre J, Relman DA, editors. MBio. 2015 Jul;6(4).

13. Prokesch BC, TeKippe M, Kim J, Raj P, TeKippe EM, Greenberg DE. Primary osteomyelitis caused by hypervirulent Klebsiella pneumoniae. Lancet Infect Dis. 2016 Sep;16(9):e190–5.

14. Patel PK, Russo TA, Karchmer AW. Hypervirulent Klebsiella pneumoniae. Open Forum Infect Dis. 2014 Jun;1(1):ofu028–ofu028.

15. Wang Q, Li B, Tsang AKL, Yi Y, Woo PCY, Liu CH. Genotypic Analysis of Klebsiella pneumoniae Isolates in a Beijing Hospital Reveals High Genetic Diversity and Clonal Population Structure of Drug-Resistant Isolates. Manganelli R, editor. PLoS One. 2013 Feb;8(2):e57091.

16. Chuang Y, Fang C, Lai S, Chang S, Wang J. Genetic Determinants of Capsular Serotype K1 of Klebsiella pneumoniae Causing Primary Pyogenic Liver Abscess. J Infect Dis. 2006 Mar;193(5):645–54.

17. Fang C-T, Chuang Y-P, Shun C-T, Chang S-C, Wang J-T. A Novel Virulence Gene in Klebsiella pneumoniae Strains Causing Primary Liver Abscess and Septic Metastatic Complications. J Exp Med. 2004 Mar;199(5):697–705.

18. Yeh K, Chang F, Fung C, Lin J, Siu LK. Serotype K1 Capsule, Rather than magA Per Se, Is Really the Virulence Factor in Klebsiella pneumoniae Strains That Cause Primary Pyogenic Liver Abscess. J Infect Dis. 2006 Aug;194(3):403–4.

19. Yu VL, Hansen DS, Ko WC, Sagnimeni A, Klugman KP, von Gottberg A, et al. Virulence Characteristics of Klebsiella and Clinical Manifestations of K. pneumoniae Bloodstream Infections. Emerg Infect Dis. 2007 Jul;13(7):986–93.

20. Yu W-L, Ko W-C, Cheng K-C, Lee H-C, Ke D-S, Lee C-C, et al. Association between rmpA and magA Genes and Clinical Syndromes Caused by Klebsiella pneumoniae in Taiwan. Clin Infect Dis. 2006 May;42(10):1351–8.

21. Russo TA, Olson R, MacDonald U, Metzger D, Maltese LM, Drake EJ, et al. Aerobactin Mediates Virulence and Accounts for Increased Siderophore Production under Iron-Limiting Conditions by Hypervirulent (Hypermucoviscous) Klebsiella pneumoniae. Infect Immun. 2014 Jun;82(6):2356–67.

22. Fox JG, Rohovsky MW. Meningitis caused by Klebsiella spp in two rhesus monkeys. J Am Vet Med Assoc. 1975 Oct;167(7):634–6.

23. Gonzalo A ME. Klebsiella pneumoniae infection in a New World nonhuman primate center. Lab Primate Newsl. 1991;30:13–20.

24. Pisharath HR, Cooper TK, Brice AK, Cianciolo RE, Pistorio AL, Wachtman LM, et al. Septicemia and peritonitis in a colony of common marmosets (Callithrix jacchus) secondary to Klebsiella pneumoniae infection. Contemp Top Lab Anim Sci. 2005 Jan;44(1):35–7.

25. Kindlovits A, Kindlovits L. Clínica e Terapêutica em Primatas Neotropicais. 2nd ed. Rio de Janeiro: L.F. Livros; 2009. 535 p.

26. Chalmers DT, Murgatroyd LB, Wadsworth PF. A survey of the pathology of marmosets (Callithrix jacchus) derived from a marmoset breeding unit. Lab Anim. 1983 Oct;17(4):270–9.

27. Richard C. [Epidemiology of Klebsiella pneumoniae infections in 2 colonies of squirrel monkeys and lemurs]. Bull Soc Pathol Exot Filiales. 1989;82(4):458–64.

28. Soto E, LaMon V, Griffin M, Keirstead N, Beierschmitt A, Palmour R. Phenotypic and genotypic characterization of Klebsiella pneumoniae isolates recovered from nonhuman primates. J Wildl Dis. 2012 Jul;48(3):603–11.

29. Twenhafel NA, Whitehouse CA, Stevens EL, Hottel HE, Foster CD, Gamble S, et al. Multisystemic Abscesses in African Green Monkeys (Chlorocebus aethiops) with Invasive Klebsiella pneumoniae—Identification of the Hypermucoviscosity Phenotype. Vet Pathol. 2008 Mar;45(2):226–31.

30. Burke RL, Whitehouse CA, Taylor JK, Selby EB. Epidemiology of invasive Klebsiella pneumoniae with hypermucoviscosity phenotype in a research colony of nonhuman primates. Comp Med. 2009 Dec;59(6):589–97.

31. Guerra MFL, Teixeira RHF, Ribeiro VL, Cunha MPV, Oliveira MGX, Davies YM, et al. Suppurative peritonitis by Klebsiella pneumoniae in captive gold-handed tamarin (Saguinus midas midas). J Med Primatol. 2016 Feb;45(1):42–6.

32. Anzai EK, de Souza Júnior JC, Peruchi AR, Fonseca JM, Gumpl EK, Pignatari ACC, et al. First case report of non-human primates (Alouatta clamitans) with the hypervirulent Klebsiella pneumoniae serotype K1 strain ST 23: A possible emerging wildlife pathogen. J Med Primatol. 2017 Dec;46(6):337–42.

33. Wu F, Ying Y, Yin M, Jiang Y, Wu C, Qian C, et al. Molecular Characterization of a Multidrug-Resistant Klebsiella pneumoniae Strain R46 Isolated from a Rabbit. Int J Genomics. 2019 Aug;2019:1–12.

34. Osman KM, Hassan HM, Orabi A, Abdelhafez AST. Phenotypic, antimicrobial susceptibility profile and virulence factors of Klebsiella pneumoniae isolated from buffalo and cow mastitic milk. Pathog Glob Health. 2014 Jun;108(4):191–9.

35. Boszczowski I, Salomão MC, Moura ML, Freire MP, Guimarães T, Cury AP, et al. Multidrug-resistant Klebsiella pneumoniae: genetic diversity, mechanisms of resistance to polymyxins and clinical outcomes in a tertiary teaching hospital in Brazil. Rev Inst Med Trop Sao Paulo. 2019;61.

36. Brasil Ministério da Saúde, Secretaria de Vigilância em Saúde, Doenças D de V das, Transmissíveis. Guia de vigilância de epizootias em primatas não humanos e entomologia aplicada à vigilância da febre amarela / Ministério da Saúde, Secretaria de Vigilância em Saúde, Departamento de Vigilância das Doenças Transmissíveis. 2nd ed. Ministério da Saúde, editor. Brasília; 2014. 100 p.

37. Hunter SB, Vauterin P, Lambert-Fair MA, Van Duyne MS, Kubota K, Graves L, et al. Establishment of a universal size standard strain for use with the PulseNet standardized pulsed-field gel electrophoresis protocols: converting the national databases to the new size standard. J Clin Microbiol [Internet]. 2005;43(3):1045–50. Available from: http://www.ncbi.nlm.nih.gov/pubmed/15750058

38. Gerlach GF, Clegg S, Allen BL. Identification and characterization of the genes encoding the type 3 and type 1 fimbrial adhesins of Klebsiella pneumoniae. J Bacteriol. 1989;171(3):1262–70.

39. Tacconelli E, Carrara E, Savoldi A, Harbarth S, Mendelson M, Monnet DL, et al. Discovery, research, and development of new antibiotics: the WHO priority list of antibiotic-resistant bacteria and tuberculosis. Lancet Infect Dis. 2018 Mar;18(3):318–27.

40. Cox BL, Schiffer H, Dagget G, Beierschmitt A, Sithole F, Lee E, et al. Resistance of Klebsiella pneumoniae to the innate immune system of African green monkeys. Vet Microbiol. 2015 Mar;176(1–2):134–42.

41. Ludlage E, Mansfield K. Clinical care and diseases of the common marmoset (Callithrix jacchus). Comp Med. 2003 Aug;53(4):369–82.

42. Antunes SG, de Groot NG, Brok H, Doxiadis G, Menezes AAL, Otting N, et al. The common marmoset: A new world primate species with limited Mhc class II variability. Proc Natl Acad Sci. 1998 Sep;95(20):11745–50.

43. Russo TA, Olson R, Fang C-T, Stoesser N, Miller M, MacDonald U, et al. Identification of Biomarkers for Differentiation of Hypervirulent Klebsiella pneumoniae from Classical K. pneumoniae. Diekema DJ, editor. J Clin Microbiol. 2018 Jun;56(9).

44. Cubero M, Grau I, Tubau F, Pallarés R, Dominguez MA, Liñares J, et al. Hypervirulent Klebsiella pneumoniae clones causing bacteraemia in adults in a teaching hospital in Barcelona, Spain (2007–2013). Clin Microbiol Infect. 2016 Feb;22(2):154–60.

45. Liu Y, Long D, Xiang T-X, Du F-L, Wei DD, Wan L-G, et al. Whole genome assembly and functional portrait of hypervirulent extensively drug-resistant NDM-1 and KPC-2 co-producing Klebsiella pneumoniae of capsular serotype K2 and ST86. J Antimicrob Chemother. 2019 May;74(5):1233–40.

46. Decre D, Verdet C, Emirian A, Le Gourrierec T, Petit J-C, Offenstadt G, et al. Emerging Severe and Fatal Infections Due to Klebsiella pneumoniae in Two University Hospitals in France. J Clin Microbiol. 2011 Aug;49(8):3012–4.

47. Bialek-Davenet S, Nicolas-Chanoine M-H, Decré D, Brisse S. Microbiological and clinical characteristics of bacteraemia caused by the hypermucoviscosity phenotype of Klebsiella pneumoniae in Korea. Epidemiol Infect. 2013 Jan;141(1):188–188.

48. Bialek-Davenet S, Criscuolo A, Ailloud F, Passet V, Nicolas-Chanoine M-H, Decre D, et al. Development of a multiplex PCR assay for identification of Klebsiella pneumoniae hypervirulent clones of capsular serotype K2. J Med Microbiol. 2014 Dec;63(Pt_12):1608–14.

49. Lin J-C, Koh T, Lee N, Fung C-P, Chang F-Y, Tsai Y-K, et al. Genotypes and virulence in serotype K2 Klebsiella pneumoniae from liver abscess and non-infectious carriers in Hong Kong, Singapore and Taiwan. Gut Pathog. 2014;6(1):21.

50. Russo TA, Gill SR. Draft Genome Sequence of the Hypervirulent Klebsiella pneumoniae Strain hvKP1, Isolated in Buffalo, New York. Genome Announc. 2013 Mar;1(2):e0006513.

51. Melot B, Brisse S, Breurec S, Passet V, Malpote E, Lamaury I, et al. Community-acquired meningitis caused by a CG86 hypervirulent Klebsiella pneumoniae strain: first case report in the Caribbean. BMC Infect Dis. 2016;16(1):736.

52. Harada S, Doi Y. Hypervirulent Klebsiella pneumoniae: a Call for Consensus Definition and International Collaboration. Diekema DJ, editor. J Clin Microbiol. 2018 Jun;56(9).

