## Supplementary material for "Detection of hypermucoviscous *Klebsiella pneumoniae* sequence type 86 capsular type K2 in South America as an unexpected cause of a fatal outbreak in captivity marmosets": Table 1

Table 1. Histologic findings described about the fragments submitted for microscopic evaluation.

| **Organ (number of analyzed specimens)** | **Histologic findings** |
| --- | --- |
| Liver (10) | Sinusoidal leukocytosis (9/10) ^1^  Hemorrhagic foci (7/10)  Necrotic foci (2/10)  Mild to moderate mononuclear cells around portal areas (3/10)  Intravascular fibrin deposition (1/10) |
| Spleen (9) | Acute necrotizing splenitis ^2^  (6/9)  Hemorrhage (8/9)  Myriads of bacilli on the red pulp (8/9) |
| Lung (10) | Subacute interstitial pneumonia (8/10)  Occasionally free bacilli (7/10)  Hemorrhage (1/10) |
| Cerebrum (10) | Nonsuppurative meningitis (2/10)  Bacilli on leptomeninges (1/10) |
| Adrenal (2) | Adrenalitis (1/2)  Foci of necrosis with perilesional bacilli and hemorrhage (1/2) |
| ^1^ Predominantly with neutrophilia (8/10)  ^2^ Acute splenitis without necrotizing pattern (2/9) | |
