## Supplementary material for "Detection of hypermucoviscous *Klebsiella pneumoniae* sequence type 86 capsular type K2 in South America as an unexpected cause of a fatal outbreak in captivity marmosets": Table 2

Table 2. Antimicrobial susceptibility profile determined by Sensititre (Thermo Scientific) of the epizootic strain hypermucoviscous *K. pneumoniae* ST86 strain P04.

| **Antimicrobial agent** | **Abbreviation** | **Minimal inhibitory concentration (mg/L)** | **Category** |
| --- | --- | --- | --- |
| Amikacin | AK | <4,0 | Susceptible |
| Ampicillin-sulbactam | A/S | 8/4 | Susceptible |
| Aztreonam | AZT | <2 | Susceptible |
| Cefepime | FEP | <2 | Susceptible |
| Cefotaxime | FOT | <2 | Susceptible |
| Ceftazidime | TAZ | <1,0 | Susceptible |
| Ciprofloxacin | CIP | <0,06 | Susceptible |
| Colistin | COL | <0,25 | Susceptible |
| Doripenem | DOR | <0,5 | Susceptible |
| Doxycycline | DOX | 2,0 | Susceptible |
| Gentamicin | GEN | <1,0 | Susceptible |
| Imipenem | IMI | <1,0 | Susceptible |
| Levofloxacin | LEVO | <1,0 | Susceptible |
| Meropenem | MERO | <1,0 | Susceptible |
| Minocycline | MIN | 4,0 | Susceptible |
| Piperacillin-tazobactam | PT | <8/4 | Susceptible |
| Polymyxin B | POL | <0,25 | Susceptible |
| Sulfamethoxazole-trimethoprim | SXT | <0,5/9,5 | Susceptible |
| Ticarcillin-Clavulanic Acid | TIM | <16/2 | Susceptible |
| Tigecycline | TGC | 0,5 | Susceptible |
| Tobramycin | TOB | <1,0 | Susceptible |
| 1. Tigecycline breakpoint followed the Food and Drug Administration (FDA) recommendations. | | | |
